# Sopa: a technology-invariant pipeline for analyses of image-based spatial-omics

**DOI:** 10.1101/2023.12.22.571863

**Authors:** Quentin Blampey, Kevin Mulder, Charles-Antoine Dutertre, Margaux Gardet, Fabrice André, Florent Ginhoux, Paul-Henry Cournède

## Abstract

Spatial-omics data allow in-depth analysis of tissue architectures, opening new opportunities for biological discovery. In particular, imaging techniques offer single-cell resolutions, providing essential insights into cellular organizations and dynamics. Yet, the complexity of such data presents analytical challenges and demands substantial computing resources. Moreover, the proliferation of diverse spatial-omics technologies, such as Xenium, MERSCOPE, CosMX in spatial-transcriptomics, and MACSima and PhenoCycler in multiplex imaging, hinders the generality of existing tools. We introduce Sopa (https://github.com/gustaveroussy/sopa), a technology-invariant, memory-efficient pipeline with a unified visualizer for all image-based spatial omics. Built upon the universal SpatialData framework, Sopa optimizes tasks like segmentation, transcript/channel aggregation, annotation, and geometric/spatial analysis. Its output includes user-friendly web reports and visualizer files, as well as comprehensive data files for in-depth analysis. Overall, Sopa represents a significant step toward unifying spatial data analysis, enabling a more comprehensive understanding of cellular interactions and tissue organization in biological systems.

## 1 Introduction

Spatial-omics data offer opportunities to improve our understanding of cellular interactions within their micro-environment and the intricacies of tissue organization^1,2^. Recent advancements in imaging technologies have expanded these capabilities, enabling the measurement of 1000+ genes through Spatial Transcriptomics^3^ and/or the analysis of 50+ proteins via Multiplex Imaging^4^. These include Merfish^5^, ISH^11^, ISS^6^, MICS^7^, PhenoCycler^8^ and IMC^9^, all of which provide single-cell resolution, previously unachieved with spot-based techniques like 10X Visium^10^ or Nanostring GeoMX^12^. Therefore, image-based technologies provide a higher resolution — up to the subcellular level — which is needed for a detailed exploration of individual cells and their gene expression profiles within their spatial context. This level of precision has been essential for unravelling tissue architecture and understanding cellular interactions; it marks the beginning of a significant leap forward in our comprehension of biological systems^8,13,14^.

In parallel with these technological advancements, the analysis of image-based spatial omics has encountered significant computational challenges and limitations^3,15–18^. Most existing methods^19–21^ are not designed to handle large images with millions of cells. Their usage typically demands high-performance computational clusters with substantial memory resources, which limits accessibility to spatial omics due to cost and hardware constraints. As a result, most companies have developed proprietary tools for their own data types, primarily focusing only on segmentation and visualization. Yet, these proprietary tools have certain constraints, such as (i) a limit on specific functionalities, (ii) no incorporation of the latest state-of-the-art methods, and (iii) a lack of versatility, as they cannot be applied to other technologies. Two other limitations related to tool diversity are that there is a learning and adaptation process for each suite and that the tools’ specificities lead to variations in the analysis of comparable data types. In a similar fashion, current open-source analysis libraries often rely on (i) already-segmented data^22,23^, (ii) specific data types^24,25^, or (iii) a subset of analysis tasks^24,25^, resulting in fragmented approaches and difficulty in adapting one approach to a different type of technology. The absence of a unified data representation and modular programming interface further complicates the integration of various analysis steps.

To address these gaps, our work introduces Spatial Omics Pipeline and Analysis, or Sopa, a novel computational framework that enhances the accessibility, efficiency, and interpretability of image-based spatial-omics data. Sopa is a memory-efficient pipeline that works across all image-based spatial-omics technologies and that can display results in a shared visualizer. This includes the most recent Spatial Transcriptomics technologies (Xenium, MERSCOPE, CosMX) and also the multiplex imaging techniques (e.g., MACSima, PhenoCycler, Hyperion). Sopa’s capabilities include segmentation and multilevel annotation, both based on transcripts and/or stainings, as well as spatial statistics and niche geometry analysis. We demonstrate Sopa’s performance on four public datasets: two spatial-transcriptomics (MERSCOPE, Xenium) and two multiplex imaging technologies (PhenoCycler, MACSima), and provide a memory and time benchmark over multiple dataset sizes. Additionally, we demonstrate Sopa’s capabilities for geometric and spatial analysis on the MERSCOPE dataset by analyzing cell colocalization with regard to cell types and niches, showing promise for biological discoveries. All these functionalities are accessible via our open-source code, which includes a Command Line Interface (CLI), an Application Programming Interface (API), and a flexible Snakemake^26^ workflow, enabling users with various levels of expertise to process spatial-omics data seamlessly, from no-code simplicity to full flexibility. The pipeline’s generic nature ensures effortless transitions to other types of spatial-omics data, making it a versatile and powerful tool for the scientific community.

## 2 Results

### 2.1 Technology-invariant pipeline

To establish versatile tools, a common strategy involves adopting a shared data structure that seamlessly integrates across diverse technologies. SpatialData^29^ serves as one such comprehensive framework, including readers tailored for the most widely used spatial-omics technologies. Building upon this, Sopa converts any data into a SpatialData object, on which all of the six following tasks are performed. First, if needed, users can interactively select a region of interest, facilitating the exclusion of less relevant or lower-quality areas. Next, we generate overlapping patches of images and/or transcripts. Segmentation can than be performed for each individual patch, and we currently support Cellpose^19^ (image-based segmentation) and Baysor^20^ (transcripts-based segmentation). Afterwards, the cell segmentation masks are converted into polygons and merged over all patches to remove potential artefacts. Following these first four steps, we average the staining intensities and count the transcripts inside each cell (see subsection 3.4 and subsection 3.5), allowing further tasks such as annotation. For example, Sopa currently supports Tangram^21^ for transcript-based annotation, and a simple Z-score method for staining-based annotation (subsection 3.7). Finally, we implemented spatial and geometric analysis tools to fully exploit the spatial nature of the data (subsection 3.8). For convenience, all image-based technologies can be visualized in a shared explorer (see subsection 2.2), and an HTML report is provided for pipeline quality checks. The full process described above is summarised in Figure 1.

**Figure 1:**
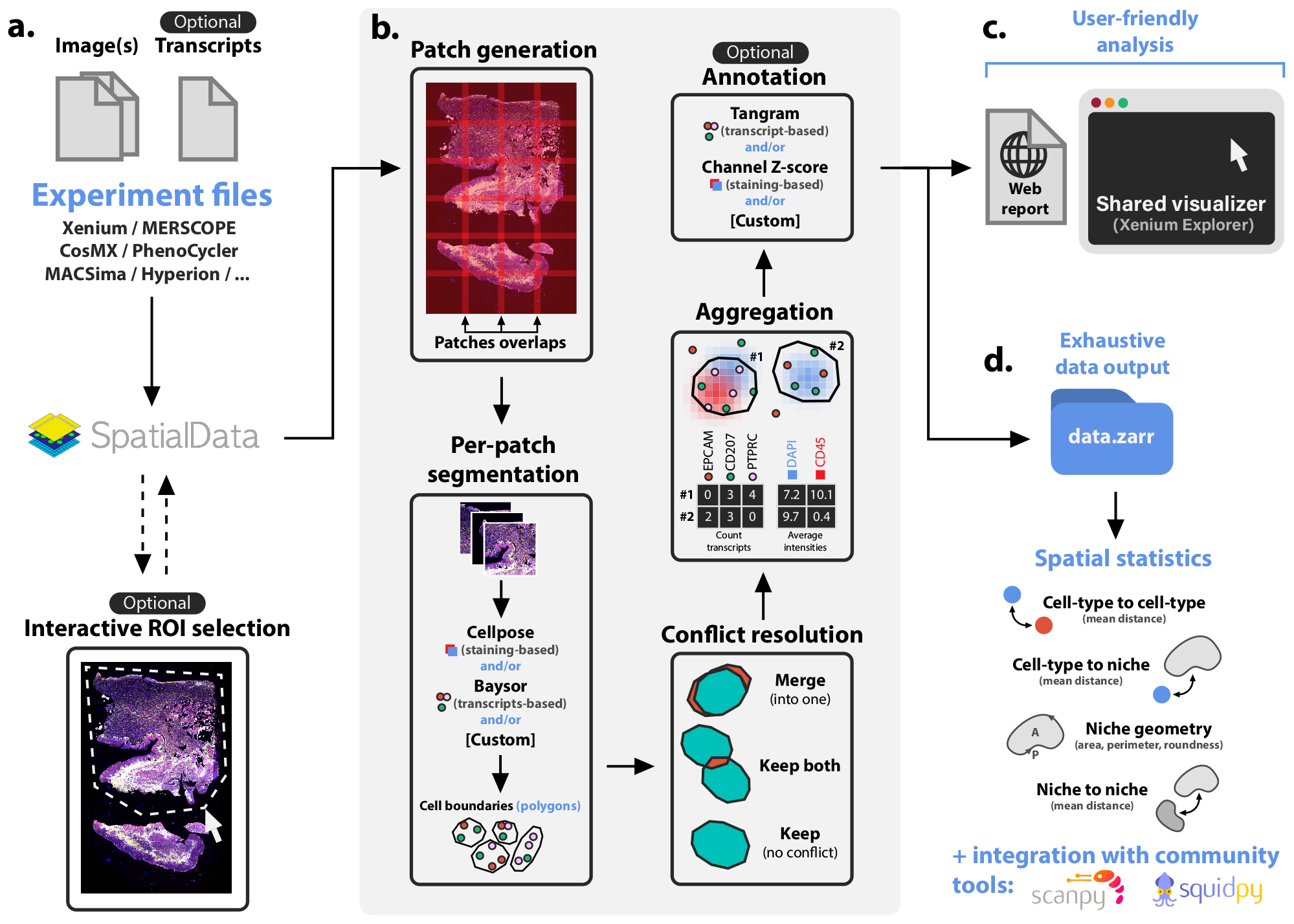
Overview of Sopa. **a**. The pipeline input consists of experimental files of any image-based spatial-omics. It is transformed into a SpatialData object, on which we can optionally select a region of interest (ROI) interactively. **b**. Afterwards, the data is split into overlapping patches, and segmentation is run on each patch (for instance, Cellpose, Baysor, or a custom segmentation tool). Since patches are overlapping, some cells can be segmented multiple times on different patches. Therefore, these conflicts have to be resolved: two boundaries with a significant overlap are merged into one cell, while two cells barely touching are kept separate. The next step is aggregation, i.e., counting transcripts and averaging each channel intensity inside each cell. This allows annotation, either based on transcripts (using Tangram) or on channel intensities. **c**. Afterwards, Sopa outputs a user-friendly report and files to be opened in the Xenium Explorer (whatever the input technology). **d**. All data files are kept for further analysis in Sopa, such as spatial statistics, or integration with community tools.

### 2.2 Shared interactive visualizer

In spatial-omics analysis, effective visualization is crucial but has presented challenges due to the size of the datasets. While open-source initiatives like Napari^37^ are emerging, they currently face limitations in handling large amounts of transcripts. Also, most companies provide technology-specific visualizers, offering limited user possibilities (see supplementary subsection 9.3). Yet, 10X Genomics has introduced the Xenium Explorer, an optimized visualizer whose file format is open, i.e., formats that can be generated for various SpatialData types. In Sopa, we have incorporated a converter that transforms the pipeline output into the input files compatible with the Xenium Explorer (see subsection 3.6 and Figure 1c). This integration ensures access to an efficient and robust visualizer, extending its functionalities to any technology whose data is readable by Sopa. Importantly, this adaptation applies to both spatial transcriptomics and multiplex imaging data, with the “Transcripts” panel selectively available for transcriptomics data. The Figure 3b/e show views using this Explorer, while supplemental Figure 1/Figure 2 are full-window examples. In addition to visualisation, the Xenium Explorer contains an interactive tool to align images from which we can export a transformation matrix and use it to align images on the SpatialData object to benefit from all the functionalities in Sopa (see supplementary subsection 9.5).

**Figure 2:**
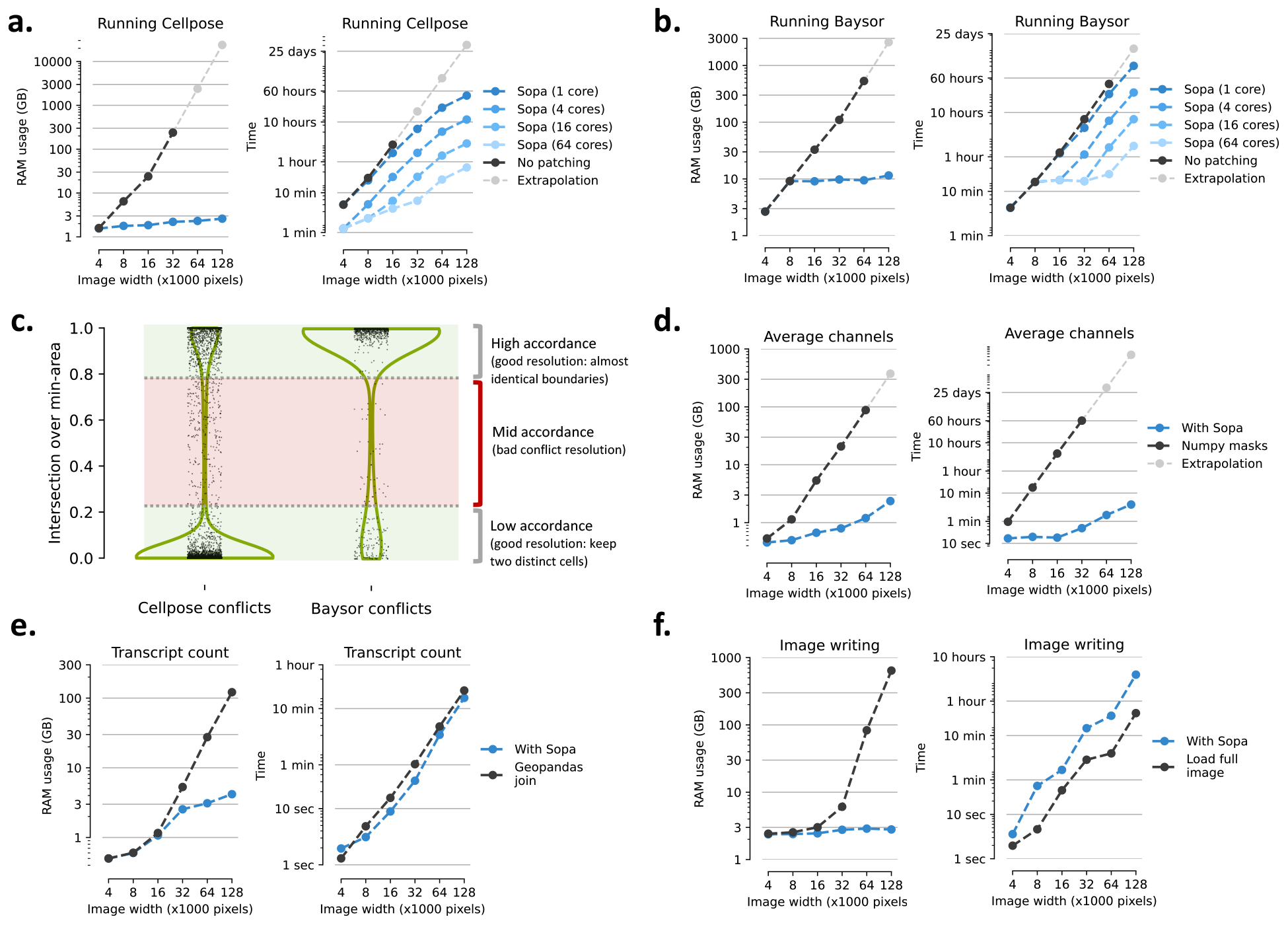
Computational efficiency of Sopa in terms of RAM and time on different dataset sizes. **a**. Cellpose segmentation comparison: with and without patching. The RAM usage is given per core. **b**. Baysor segmentation comparison: with and without patching. The RAM usage is given per core. **c**. Violin plots showing the intersection-over-min-area density of segmentation conflicts when using patches (for both Cellpose and Baysor). When resolving a conflict, the two good cases are either (i) a high concordance between the two cells (which will be merged), or (ii) a low concordance between them (the two cells are kept). Anything between 0.2 and 0.8 is considered a bad segmentation overlap and could deteriorate further analyses. **d**. Channels averaging for each cell: Sopa and standard average inside numpy masks. **e**. Counting each gene inside each cell: with Sopa compared to GeoPandas join operation on the whole DataFrame. **f**. Writing image as a tiff file for the Xenium Explorer: with Sopa compared to what is recommended by 10X Genomics, i.e. loading the whole image in memory.

**Figure 3:**
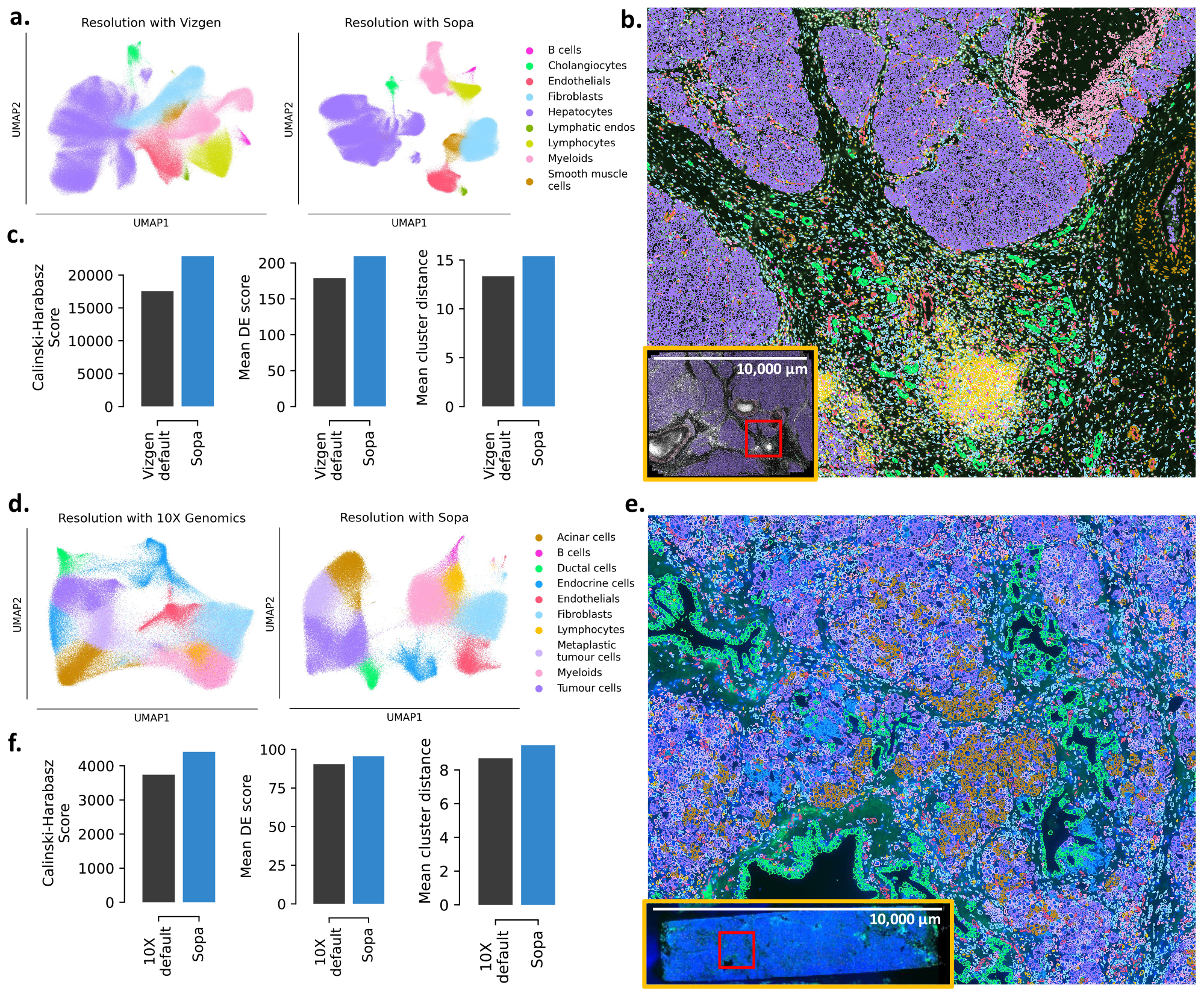
Data resolution after Sopa segmentation compared to proprietary segmentation over two spatial-transcriptomics technologies: MERSCOPE (a-c) and Xenium (d-f). **a**. UMAPs of cells after Vizgen default segmentation on the MERSCOPE dataset (left) and after Sopa segmentation on the same dataset (right). **b**. Visualization of cell types on the MERSCOPE dataset after annotation with Sopa. Colours correspond to the legend of (a). **c**. Three cluster separation metrics compare the quality of these two segmentations on the MERSCOPE dataset. **d**. UMAPs of cells after 10X Genomics default segmentation on the Xenium dataset (left) and after Sopa segmentation on the same dataset (right). **e**. Visualization of cell types on the Xenium dataset after annotation with Sopa. Colours correspond to the legend of (d). **f**. Three cluster separation metrics compare the quality of these two segmentations on the Xenium dataset.

### 2.3 Memory efficiency of Sopa

Managing large datasets is a critical challenge in spatial omics, particularly when dealing with images that can reach hundreds of gigabytes and contain hundreds of millions of transcripts in spatial transcriptomics data. This necessitates implementing memory optimization techniques to ensure the scalability of the analysis. Notably, segmentation algorithms like Cellpose^19^ and Baysor^20^ encounter scalability issues with large images, as illustrated in Figure 2a/b. To tackle this, these segmentation models are applied to smaller regions called patches, drastically decreasing random-access-memory (RAM) usage and time. While this patching process generates possible segmentation conflicts, we show in Figure 2c that this does not impact segmentation quality, since few conflicts are difficult to resolve, i.e., with a score between 0.2 and 0.8. Additionally, the conventional storage of cell boundaries as raster masks demands significant memory for storage and processing (see Figure 2d). To address this, we adopt a more efficient approach by storing cell boundaries as polygons using Shapely^30^, which proves highly effective for both on-disk and in-memory storage. This also facilitates geometry-related operations, such as cell expansion, area/perimeter computations, and cell-cell intersections. Combined with the image lazy loading offered by SpatialData^29^ and Xarray^34^, we implement a fast channel averaging on cell boundaries by combining geometry operations and image chunk lazy loading (see Figure 2d), i.e., deferring loading until needed for processing. Additionally, using memory-efficient tools like Dask^31^, we extend geometric operations of GeoPandas^32^ on chunks of transcripts, ensuring parallel processing of as many chunks as possible without exceeding memory limits (see Figure 2e). For image conversion to a pyramidal ‘.tif’, we significantly lower the memory usage compared to what is recommended by 10X Genomics (see subsection 3.6) by writing tiles in a lazy manner, which avoids loading the full image in memory (see Figure 2f). To highlight Sopa’s memory efficiency, we compared its RAM usage against standard methods for all tasks mentioned above across various dataset sizes, summarized in Figure 2. Overall, the latter figure shows significant improvements in terms of RAM and time: depending on the tasks, Sopa can require between 10 and 100 times less memory than normal techniques and can be up to 100 times faster. Even on the largest image, Sopa can be run with a simple laptop with 16GB of RAM.

### 2.4 Wide use cases and customization

Sopa offers three distinct options, each tailored to different use cases: (i) a Snakemake^26^ pipeline that enables a quick start within minutes, (ii) a CLI that facilitates rapid prototyping of a personalized pipeline, and (iii) an API that allows direct usage of Sopa as a Python package^1^, providing full flexibility and customization. The Snakemake pipeline remains consistent across various technologies, with only its configuration differing. Users can leverage existing configuration files, selecting one that aligns with their technology, which then enables them to execute the pipeline without any code updates. Another advantage of Sopa’s generality and scalability is that more advanced users seeking customisable pipelines can use the CLI. Notably, Sopa’s general design allows for an easy integration of new or custom segmentation methods, rendering them memory-efficient and accessible for all image-based spatial-omics applications. Additionally, the Python API is available for users interested in incorporating specific parts of Sopa into their personal libraries. This API also facilitates integration with other tools of the scverse^44^ ecosystem, such as Scanpy^33^ or Squidpy^22^ (see supplemental subsection 9.2). In particular, the integration with Squidpy enables the use of post-processing tools for cell-cell interaction and spatially variable gene analysis.

### 2.5 High resolution of the tumour microenvironment

Segmentation plays a crucial role in image-based spatial-omics analysis. Sopa focuses significantly on improving this step (see subsection 3.3) by enabling the usage of state-of-the-art segmentation models like Baysor^20^ on large datasets. Indeed, as shown on Figure 2a/b, these high-quality segmentation tools use a lot of memory, which hinders their usage on large spatial datasets. To evaluate the resolution provided by Sopa after segmentation, we annotated major cell types and conducted tests on four datasets: two spatial-transcriptomics datasets (MERSCOPE and Xenium) and two multiplex-imaging datasets (PhenoCycler and MACSima)^2^. For the MERSCOPE and Xenium datasets, proprietary segmentations were provided by Vizgen and 10X Genomics, respectively. In comparison to these segmentations, Sopa shows an improved cell-type distinction on UMAP^28^ plots (see Figure 3a/d) by leveraging Baysor. To support these visual observations, we used multiple metrics (see subsection 3.2), indicating that Sopa can generate more significant population-specific genes, greater intra-cluster distance, and improved cluster separation (see Figure 3c/f). The increased resolution in spatial omics data allows for a more in-depth exploration compared to previous segmentations (see supplementary Figure 3 for more details).

Sopa also facilitates the concurrent analysis of both RNA and proteins. To demonstrate this, we used the Xenium dataset, which includes transcriptomic expression and protein stainings (CD20, PPY and TROP2). CD20 is a common marker for B cells, PPY is expressed by endocrine cells, and TROP2 is overexpressed in tumour cells. 10X Genomics currently does not produce files with protein expression per cell, while Sopa does support the analysis of proteins. To demonstrate, we aligned the Xenium staining image to the original coordinate system (see supplemental subsection 9.5), and Sopa computed the CD20/PPY/TROP2 intensity within all cell boundaries. Combined with transcriptomic expression, CD20 staining greatly facilitates the annotation of B cells, as shown by their clear delimitation on Figure 3d and Figure 3c. In the future, we expect technologies to be able to run more protein stainings in parallel with transcriptomics data, making this kind of analysis even more valuable.

Regarding multiplex imaging, Sopa shows efficiency in (i) managing large protein staining panels and (ii) segmenting millions of cells (using Cellpose). The former is exemplified by the MACSima dataset with 61 stained proteins. Again, we computed staining intensity per cell, and Figure 4a demonstrates Sopa’s capacity to annotate high-resolution cell types. Secondly, the PhenoCycler dataset underscores Sopa’s ability to handle datasets of substantial size, with an area of 3 squared centimetres, containing approximately 2,500,000 cells. The corresponding cell resolution is shown in Figure 4c.

**Figure 4:**
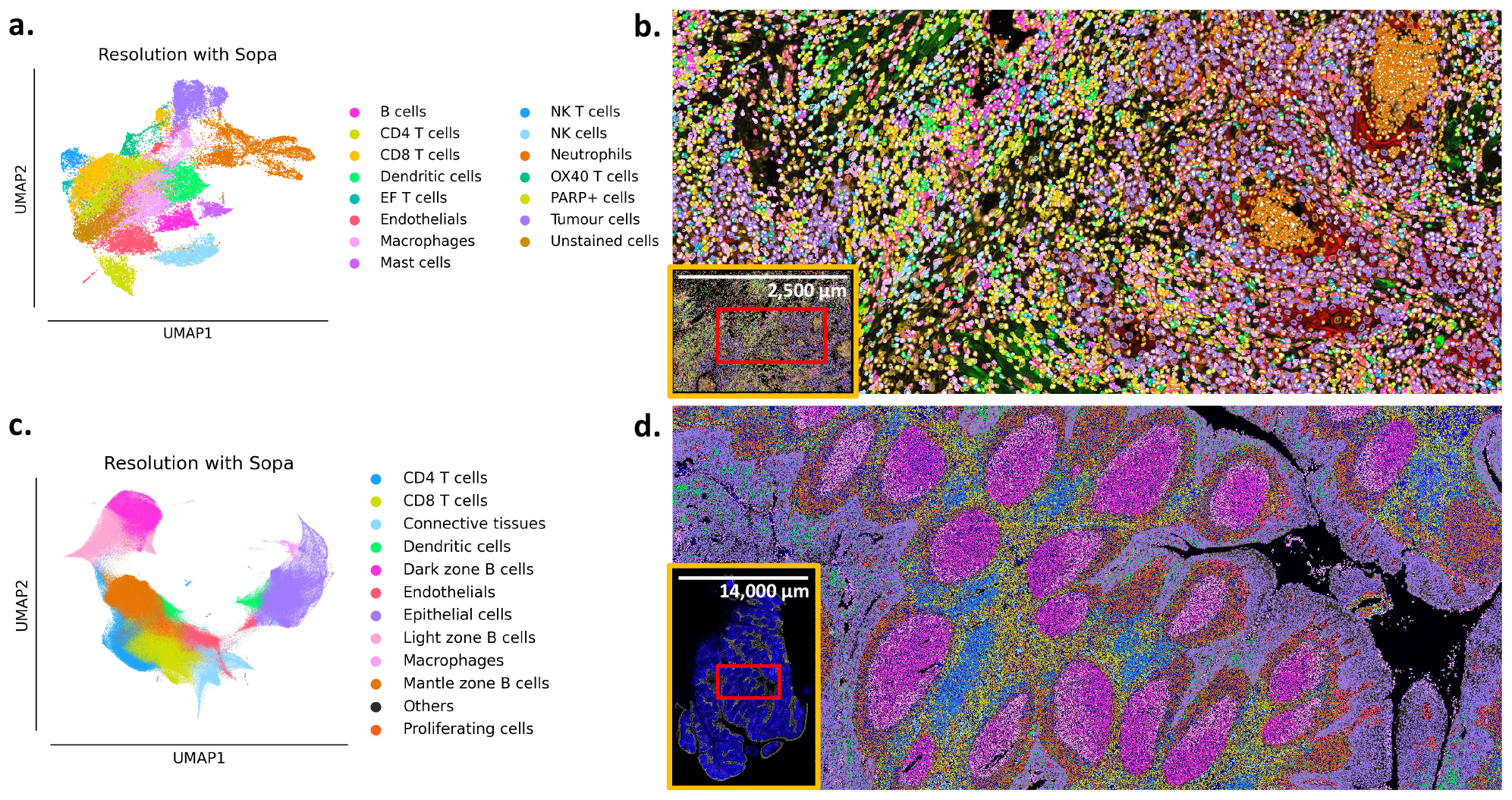
Data resolution after Sopa segmentation over two multiplex-imaging technologies: MACSima (a-b), and PhenoCycler (c-d). **a**. UMAP representing cell types on the MACSima dataset, annotated based on 61 protein stainings. **b**. Cells of the MACSima dataset visualized. The colours correspond to the legend of (a). **c**. UMAP representing cell types on the PhenoCycler dataset, annotated based on 31 protein stainings. This represents approximately 2,500,000 cells. **d**. Cells of the PhenoCycler dataset visualized. The colours correspond to the legend of (c).

In summary, these studies demonstrate that Sopa can (i) be applied across diverse technologies, (ii) efficiently handle millions of cells, and (iii) seamlessly operate on both transcriptomics and protein stainings.

### 2.6 Geometric and spatial analyses

Spatial omics naturally unlock multiple biological questions related to spatial organization. While some are addressed in libraries such as Squidpy^22^, metrics related to the distance between cell-types/niches and the geometric characteristics of those niches are not provided. These metrics could help in the understanding of the morphology of the tumour micro-environment and its location with regards to different cell types. Such statistics have been shown to be relevant for predicting disease progression or response to treatment^54,55^. For instance, it is known that tertiary lymphoid structures (TLS) have a good prognosis^47^, but their geometry has not been studied. TLS may come in different sizes, shapes, occurrences, or locations with regard to other niches. Such statistics are generalized in subsection 3.8 for all cell categories (usually, cell types or niches). Leveraging this spatial analysis, we demonstrate a better understanding of the dynamics among different cell types and their relation to different spatial niches on the MERSCOPE liver dataset (Figure 5). To use Sopa geometric analysis, we run STAGATE^27^ to identify eight distinct niches (or “spatial domains”) across various tumour regions (Figure 5a). First, we show in Figure 5b four geometric properties related to these niches: for each niche compartment, we counted their occurrence on the same slide, as well as their mean area, perimeter, and roundness (see subsection 3.8 for more details). For instance, our geometric analysis shows a high occurrence of vascular niches, that are small in area and perimeter, but have a high roundness. Conversely, the stroma has only one occurrence and is highly “unround”, and Figure 5c shows that this shape enables a “proximity” to every other niche. Figure 5c also highlights how far the vascular niche is from the necrosis. While such observations are not novel, our geometric computation allows for statistical comparisons over multiple patients, which could lead to the discovery of significant geometric biomarkers in large-scale studies.

**Figure 5:**
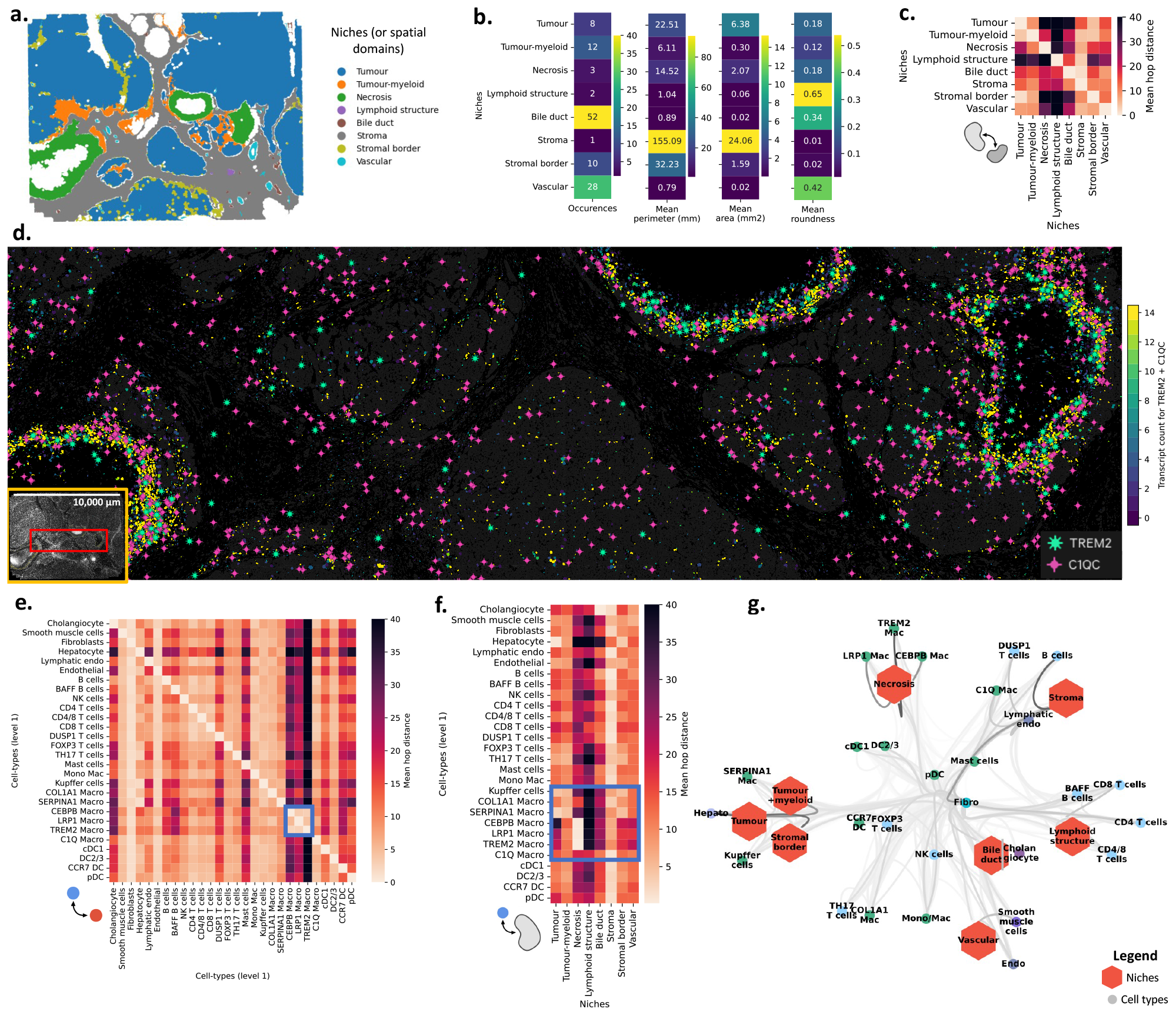
Geometric analyses and spatial statistics on the MERSCOPE liver dataset. **a**. Niches (or spatial domains) after geometric conversion to shapely polygons. **b**. Geometric statistics of the niches: their occurrence, perimeter, area, and roundness. **c**. Heatmap of average hop distance between niches and niches. **d**. Localisation of TREM2 macrophages shown in the visualizer. The TREM2 and C1QC genes are shown, and cells are coloured by their gene counts for the two selected genes. **e**. Heatmap of average hop distance between cell types and all other cell types. LRP1, CEBP, and TREM2 macrophages show a high proximity. **f**. Heatmap of average hop distance between cell types and niches. The macrophage subpopulations show heterogeneous localisation with respect to the niches. LRP1, CEBP, and TREM2 macrophages are enriched in the necrosis niche. **g**. Network plot summarising the distance metrics of (c)/(e)/(f). Each node of the network corresponds either to a niche (hexagon) or a cell type (circle). The lower the mean distance between the two nodes, the higher the weight of the edge between these two nodes. A high node-node proximity is shown by a dark edge. Overall, it provides an overview of the colocalisation of cell types and niches in the tumour environment.

We also utilised Sopa to assess the intricacies of the tumour complexity. We annotated the immune populations of the MERSCOPE dataset in higher definition (see supplementary Figure 4a/b) and in parallel performed a differential analysis on each niche to better understand niche complexity. This revealed a distinct necrotic niche correlated with TREM2 macrophages (expressing TREM2, C1QC and CSF1R), a population of macrophages reported across cancer types and often associated with bad prognosis^50,51^ (see Figure 5d and Figure 4c). To deepen this understanding of tissue intricacies, we investigated whether these TREM2 macrophages were in close distance with any other cell type (see Figure 5e). Strikingly, this figure highlighted that three macrophage populations (LRP1, CEBP, and TREM2-macrophages) exclusively interacted with themselves. Correlating their location with the niche revealed that their co-occurrence is specific to the necrotic niche (see Figure 5f). When combining all (cell-cell/cell-niche/niche-niche) interactions, this affirms again the association of LRP1/CEBP/TREM2-macrophages in the necrotic niche, yet it also highlights the heterogeneity of all macrophage populations and their relation to the niche in the whole tissue environment (see Figure 5g). These combined interactions also showed that, inversely, the conventional dendritic cells (DCs) are not associated with any niche environment, accentuating how some populations can also be niche-independent. This observed spatial location underscores a potential reprogramming feature of macrophages based on their specific niche. While it is known that the accumulation of TREM2 macrophages has been associated with enrichment in the tumour regions^45,48,49^, Sopa can provide insights for a refined understanding of a macrophage-specific tumour-associated phenotype. These examples illustrate that this geometric and spatial analysis — computed with Sopa — helps better understand the tumour’s architecture and its relationship with cell type phenotypes.

## 3 Online Methods

### 3.1 Datasets used

Four public datasets were used to demonstrate Sopa’s abilities. First, we used a MERSCOPE dataset (from Vizgen) of the human liver Hepatocellular carcinoma (HCC), called FFPE Human Immuno-oncology Data Set May 2022. It is composed of a 500-gene panel, and has DAPI staining and PolyT staining. It contains about 500,000 cells, depending on the segmentation. Secondly, we used a Xenium dataset (from 10X Genomics) of pancreatic cancer (adenocarcinoma, Grade I-II) with the Xenium Human Multi-Tissue and Cancer Panel, in parallel with corresponding H&E image, and a protein-staining image with DAPI/CD20/PPY/TROP2. Note that the two latter images has to be aligned on the default main DAPI image. It contains about 180,000 cells, depending on the segmentation. Thirdly, we used a PhenoCycler dataset (from Akoya Biosciences) of the human tonsil (FFPE) with 31 protein stainings. It contains about 2,500,000 cells, depending on the segmentation. Finally, we used a MACSima dataset (from Miltenyi) of head and neck squamous cell carcinoma (HNSCC) with 61 protein stainings. It contains about 40,000 cells, depending on the segmentation. For more details about the accessibility of these datasets, see section 7.

### 3.2 Metrics used and computational details

The Calinski-Harabasz-Score is defined as the ratio of the sum of between-cluster dispersion and of within-cluster dispersion. To compute this score, we used the implementation in scikit-learn^40^. The mean cluster distance is the average distance between all pairwise combinations of cells between two different clusters; thus, a higher distance indicates a better cluster separation. For the differential expression analysis, we ran the scanpy^33^ *rank_genes_groups* function, and we averaged the score of the 20 most significant genes for each cell type. The time and memory benchmarks were performed on a Slurm cluster on the same CPU nodes. The benchmark related to Cellpose was performed on crops of the MERSCOPE dataset, while the other time and memory benchmarks were performed on a synthetic dataset (see subsection 9.6). Figure 2c was generated based on the corresponding 16,000-pixels-wide datasets; this involves 25 Cellpose patches and 4 Baysor patches. The percentage of conflicts for Cellpose (compared to all pairs of cells) was 0.007%, while this percentage was 0.001% for Baysor. The UMAPs of Figure 3 and Figure 4 were generated with scanpy^33^, using the default parameters. The MERSCOPE and Xenium datasets have been segmented with Baysor, while the PhenoCycler and MACSima datasets have been segmented with Cellpose. Both the MERSCOPE and Xenium datasets have been annotated using Tangram (see supplementary subsection 9.7 for more details).

### 3.3 Segmentation on patches

For computational efficiency, segmentation is performed on patches, i.e., small image regions. These patches have a certain overlap, which is typically chosen to be at least twice as big as the average diameter of cells (e.g., 20 microns). This way, each cell should be complete in at least one patch, which avoids artefacts due to cutting cells at the border of the patches. Subsequently, any segmentation algorithm compatible with images and/or transcripts can be applied. While Cellpose^19^ and/or Baysor^20^ are commonly used, Sopa does allow the integration of other segmentation algorithms. Following segmentation on individual tiles, the cell boundaries are transformed into polygons using Shapely^30^. Since patches overlap, some cells may be segmented across different patches, leading to segmentation conflicts where multiple polygons correspond to a single cell. To resolve this, we adopt a method similar to the one used in Vizgen’s preprocessing tool (VPT^3^). Specifically, we merge pairs of cells when the intersection area exceeds half the area of the smaller cell, ensuring a substantial overlap. If the intersection area is too small, indicating distinct cells, both polygons are retained. When the overlap area divided by the smallest cell area is close to 1, this corresponds to two almost identical cells, while a score close to 0 corresponds to two cells barely touching. On Figure 2c, we studied the distribution of this score, showing that most of the conflicts are associated with a score that is either very close to 0 or very close to 1, indicating a good conflict resolution. Additionally, note that, before segmentation, the user can decide to select a region of interest: this can be done interactively with matplotlib^36^ on a low-resolution image.

### 3.4 Channel averaging

When dealing with image-based technologies, a crucial step involves averaging the intensity of each channel within each cell. While this task can be achieved using cell masks, it proves highly inefficient in terms of both time and memory consumption. To address this challenge, we adopt a chunk-level approach: (i) For each chunk, we identify cell boundaries (i.e., polygons) that intersect with the chunk coordinates, then (ii) we determine the bounding box for each of these cells, then (iii) we extract the image values for each of these bounding boxes, and finally (iv) we rasterize the cell polygons to average the staining intensity over the local bounding box. In this way, we only load small arrays corresponding to each cell, instead of loading large cell masks. This process is repeated over all chunks, and we make sure that the channel intensity for cells located on multiple chunks is computed correctly.

### 3.5 Counting transcripts

GeoPandas^32^ is a Python library that enhances Pandas^35^ Dataframes by incorporating support for Shapely^30^ geometries. It facilitates scaling operations on geometries, making it particularly suitable for transcript counting, where transcripts can be represented as Shapely points and cells as Shapely polygons. However, the memory requirements for such operations can be substantial, especially for spatial transcriptomics datasets that may contain up to one billion transcripts. To optimize this process, we leverage Dask and execute the GeoPandas^32^ “join” operation at the partition level to assign each point (i.e., a transcript) to a polygon (i.e., a cell). Thus, each operation is carried out on smaller data frames, each less than 100MB in size. Dask efficiently assigns each partition to different workers in parallel, mitigating memory concerns. This approach proves highly effective, especially when utilizing a high-performance cluster, as Dask is designed to seamlessly scale these processes on clusters without necessitating any code modifications.

### 3.6 Conversion to the Xenium Explorer

Converting a spatial-omics object into the Xenium Explorer requires the creation of six files: (i) the image, (ii) a JSON metadata file, (iii) the cell boundaries, (iv) the cell categories (e.g., cell type or clustering), (v) the gene counts table, and (vi) the transcripts (if they exist). The conversion is done automatically by Sopa, but it can also be done manually via our CLI: `sopa explorer write <sdata_path> <output_path>`.

For image creation, a Python function is recommended in the Xenium Explorer documentation^4^ but is not optimized for large images. We updated it to support Dask^31^ arrays, i.e. (the image type used by Sopa). Pyramids of resolutions are generated via the SpatialData library^29^. To decrease memory usage, each (1024x1024) image tile is generated using an iterator that only computes the minimally required data from the Dask array at each tile generation. For higher pyramidal levels, where the image size decreases, we allow loading an image into memory if it fits, accelerating conversion.

As transcripts typically cannot be loaded entirely into memory, the Xenium Explorer avoids loading all transcripts. On low-resolution levels, only a subset of transcripts is displayed (subsampled), while zooming in reveals all transcripts from the current field of view. This pyramidal transcript view ensures low memory usage during visualization. The highest-resolution tiles are 250-micron-wide squares. For each pyramid level, the tile width doubles, and only one-fourth of the transcripts from the previous level are retained. The process stops when there is only one remaining tile that is larger than the original slide. Transcript coordinates are stored as separate chunks for each tile and resolution, saved as a Zarr file^5^. This allows loading only the transcripts corresponding to the displayed tiles when zooming in.

Cell boundaries are padded to have the same number of vertices (13). Polygon simplification is applied to polygons with more than 13 vertices using the Shapely library, reducing the number of vertices while preserving shape geometry. A fixed number of vertices enables lighter cell-boundary storage and faster visualization.

Transcript counts (cell-by-gene table) use sparse array storage. One 1D array stores all non-zero transcript counts, another array stores the cell index for each count, and a third array is a pointer indicating the gene index for these counts. Cell categories are similarly saved using indices and corresponding pointers. Once again, the file format employed is a Zarr file.

### 3.7 Cell-type annotation

#### Transcript-based annotation

Tangram^21^ is used for cell-type annotation based on an annotated scRNAseq reference. To make Tangram^21^ scalable for large datasets, we adopt a strategy of splitting the data into “bags of cells”, with the size determined by the user. This approach ensures that each Tangram iteration operates within manageable memory limits, and we subsequently merge the results to obtain the annotation for the entire dataset. Following this, Leiden^38^ clustering can be applied to refine the annotation, associating each Leiden cluster with its most prevalent Tangram cell type. Additionally, we have implemented a multi-level annotation feature based on Tangram to enhance the annotation of minor cell types if needed. The process involves initially annotating global cell populations, followed by running Tangram on specific cell groups (e.g., Myeloid cells) for a more detailed annotation (e.g., pDCs, TREM2 macrophages, etc.). All that is required is to provide multiple cell-type annotation columns in the reference scRNAseq data, and Sopa will seamlessly execute the multi-level annotation.

#### Staining-based annotation

For non-transcriptomics data, we also provide a fluorescence-based annotation. As each channel intensity is averaged inside each cell, we obtain a matrix ***X*** of shape (*N, P*), where *N* is the number of cells, and *P* the number of stainings/channels. Then, these intensities are preprocessed as in a recent article^42^:

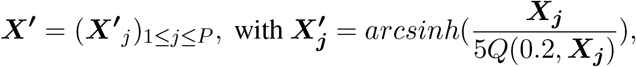

where ***X***^***′***^ is the preprocessed matrix, *arcsinh* is the inverse hyperbolic sinus function, and *Q*(0.2, ***X***_***j***_) is the 20th percentile of ***X***_***j***_. Afterwards, we use a list of stainings corresponding to a population, and each cell is annotated according to the channel whose preprocessed intensity is the highest. If desired, Leiden clustering^38^ can be run to have a deeper annotation. Each cluster can be annotated via differential analysis or by showing a heatmap of staining expression per cluster.

### 3.8 Spatial statistics

All spatial statistics are performed after computing a Delaunay graph based on the spatial location of cells. This is done with Squidpy^22^, which is itself based on Scipy^39^. We also prune long edges that cannot correspond to a physical cell-cell interaction (typically, edges longer than 40 microns). In the paragraphs below, *N* denotes the number of cells.

#### Cell category to cell-category statistics

One relevant spatial statistic is the computation of the mean or minimum distance between two cell categories. This includes the pairwise distance between cell types (e.g., the mean distance between CD8 T cells and tumour cells), as well as the distance between cell types and niches (e.g., the distance between tumour cells and tertiary lymphoid structures). Let (*C*_1_, …, *C*_*N*_) represent categories assigned to the *N* cells (e.g., cell types), and 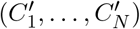 represent other categories (such as the niche to which the cell belongs). For instance, if cell “*i*” is a T cell inside the stroma, then *C*_*i*_ = “T cell” and 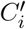 = “stroma”. The sets of unique categories are denoted *G* and *G*^*′*^, respectively; for instance, *G* can be the set of unique cell types, and *G*^*′*^ can be the set of unique niches. Then, ∀*g* ∈ *G* and ∀*g*^*′*^ ∈ *G*^*′*^, we define the mean distance between the category *g* and *g*^*′*^ as follow:

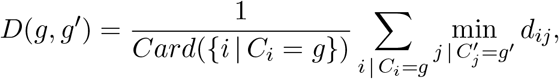

where *Card* represents the cardinal, and *d*_*ij*_ is the hop-distance between cell *i* and cell *j*. Note that 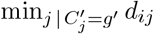 is the distance between cell *i* and the closest cell of category *g*^*′*^, that is how many hops are needed for cell *i* to “find” the category of interest. In practice, we compute *D*(*g, g*^*′*^) by multi-node graph traversal, starting from all nodes whose category is *g*^*′*^. In this way, for each *g*^*′*^ ∈ *G*^*′*^, we compute 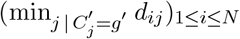 in a single graph traversal. All the resulting distances can be stored in a matrix ((*D*(*g, g*^*′*^)))_*g*∈*G,g′*∈*G′*_ and shown as a heatmap. Additionally, we combine the four matrices of distances (cell-type to cell-type, cell-type to niches, niches to cell type, and niches to niches) into an adjacency matrix whose weights are the inverse of the distance. Then, the corresponding network can be plotted using the netgraph^41^ library, as in Figure 5g, providing an interpretable visualization of the tumour microenvironment’s structure.

#### Niche geometry statistics

When niches (or spatial domains) are performed with an algorithm such as STAGATE^27^, users can decide to extract these niches as geometries to compute some relevant statistics, such as their area, perimeter, or roundness. From now on, for each cell *i*, 1 ≤ *i* ≤ *N, C*_*i*_ denotes the niche to which the cell belongs, and *G* is the corresponding set of unique niches (i.e., for all cell *i, C*_*i*_ ∈ *G*). First, we prune all the edges (*i, j*) that are in between niches from the Delaunay graph, i.e., if *C*_*i*_ ≠ *C*_*j*_. Then, we extract the connected components of the graph. Because of the way we pruned the edges, each component corresponds to one niche, but one niche can be composed of multiple components (or occurrences). For each component, we search simplices (i.e., triangles from the Delaunay graph) at the component’s border, that is, the simplices that have one or two simplex neighbours. From all the border simplices, we extract the corresponding border edges; these edges are then linked to make one or multiple rings (i.e. cyclic lines). If we have only one ring, it is transformed into a polygon, which corresponds to a “full” component. If there are multiple rings, the largest ring is the outer polygon, and the others correspond to “holes” inside the main polygon: this can happen when some components are completely surrounded by another niche. Repeating this process for all components allows the transformation of each niche *g* ∈ *G* into multiple polygons. We can then count how many occurrences (or polygons) each niche is made of, and we can also compute the mean area *A*_*g*_, perimeter *L*_*g*_, and roundness *R*_*g*_ of each niche using Shapely^30^. Note that 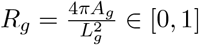, where higher values correspond to a “circle-like” shape. Also, for each niche, we filter out components whose areas are less than 5% of the area of the same niche’s largest component, as they usually correspond to low-quality artefacts from the clustering of niches.

## 4 Discussion

Advances in technology development for spatial omics hold great promise for biological discoveries. Yet, to build strong and unified foundations for spatial omics data analysis, new tools are required. With this purpose in mind, we designed and built Sopa to address several crucial aspects of spatial omics analysis: versatility, reproducibility, and scalability. It offers a suite of tools — or building blocks — designed for spatial omics, which are assembled to build a pipeline for any image-based spatial omics technology. At the end of the pipeline, it produces standardized outputs, which ease exploration and visualization. While each company’s technology comes with its own suite of tools — which differ in terms of capabilities and functionalities — Sopa does not require learning from multiple data types and software. In addition, Sopa is scalable from simple laptops to high-performance clusters, offering an extension of versatility for users.

Moreover, Sopa can easily integrate new methods and tools: as new segmentation or annotation methods are developed, they can be added to Sopa once published and validated. This incorporation into Sopa enables scalability and availability to any new technology with only minor configuration changes. As datasets become increasingly bigger, Sopa’s scalability is crucial. For instance, Sopa enabled the possibility of running Baysor on data produced by the MERSCOPE, which was previously impossible due to RAM usage and time. We also demonstrated that it significantly increases data quality compared to the default Vizgen and 10X Genomics segmentation tools. As shown on the MERSCOPE liver dataset, we were able to annotate spatial-specific macrophages, particularly TREM2 macrophages, in the necrotic niche. Additionally, TREM2 has been shown to increase with HCC, suggesting a potential immunosuppressive role of TREM2^45,49^, while necrosis has been associated with worse prognosis^52,53^. With the help of Sopa, the exploration of this relationship between tissue architecture and cell phenotypes can advance biological knowledge.

Besides higher data resolution, Sopa can also incorporate protein information into spatial analysis. Without this information, extracting the B cell population in the Xenium data would not have been possible. While current spatial technologies involve either a high number of proteins or transcripts, new developments could add extra layers of information, contributing to a better understanding of biological systems. This paper has demonstrated through various techniques that Sopa is ready to handle large multi-modal spatial technologies.

## 5 Acknowledgement

This work is supported by Prism – National Precision Medicine Center in Oncology funded by the France 2030 programme, the French National Research Agency (ANR) under grant number ANR-18-IBHU-0002, ARC Foundation and Fondation Gustave Roussy.

## 6 Code availability

The code developed in this article is available as an open-source Python package, accessible on Github at https://github.com/gustaveroussy/sopa. The code used to run the benchmark is available at https://github.com/quentinblampey/sopa_benchmark.

## 7 Data availability

The MERSCOPE dataset is freely available online at https://info.vizgen.com/merscope-ffpe-solution, and the Xenium at https://www.10xgenomics.com/resources/datasets/pancreatic-cancer-with-xenium-human-multi-tissue-and-cancer-panel-1-standard The PhenoCycler dataset is available upon request to Akoya Biosciences, see https://www.akoyabio.com/fusion/data-gallery/. The MACSima dataset is available upon request to Miltenyi.

## 9 Supplementary information

### 9.1 Choice of SpatialData as a data structure

SpatialData^29^ is a data structure developed in Python that aims to store spatial-related objects. It also provides transformations between coordinate systems (for instance, between microns and pixels), lazy representation for large images with Dask^31^ and Xarray^34^, transcripts stored as Dask^31^ dataframes, and cells polygons stored as GeoPandas^32^ polygons. The general structure of this data, the community support, and integration with the scverse^44^ ecosystem make it a reliable tool to store spatial-omics objects in Sopa. Notably, the usage of Python is appreciated since most recent models in spatial-omics are gradually moving to Python for package development^3^.

### 9.2 Integration with the scverse ecosystem

The scverse^44^ ecosystem is a Python-based suite of fundamental tools for single-cell omics data analysis. This includes the data structures SpatialData^29^ that we use for Sopa, as well as Scanpy^33^ which covers a wide range of use cases in single-cell analysis. Also, still in the scverse ecosystem, Squidpy^22^ is a Python library for the analysis of spatial single-cell data such as spatial neighbourhood analysis or ligand-receptor interaction analysis. Since Squidpy supports SpatialData, Sopa is also naturally integrating with Squidpy. Indeed, the pipeline output being a SpatialData object, Squidpy can operate on this, enabling all Squidpy functionalities to be leveraged after Sopa, or inside the pipeline. Squidpy is complementary to Sopa since it operates on processed spatial omics, contrary to Sopa, which analyses raw data. Also, the spatial statistics tools available in Sopa do not exist in Squidpy. Thus, these packages have non-overlapping and complementary functionalities.

### 9.3 Limitation of the proprietary visualization software

All visualizers are exclusive to their data structure, and require an investment of time to the users for learning their proprietary software. Besides this, some of the software comes only with the purchased machine and requires a license key for use. This limits the number of users who have a collaborative engagement and are not in possession of the machine. Data analysis from the MERSCOPE comes with a dedicated visualizer, called the “Merscope Visualizer”. Its input is proprietary *“*.*vzg”* files, a non-open format. While VPT offers the possibility to update it, a new *vzg* cannot be recreated for another type of technology. In addition, the update of this file requires performing again all required operations, even for minor changes, because everything is included in one file. Therefore, minor modifications still imply a significant runtime to be updated in the visualizer. Concerning CosMX data, they offer an online suite of tools, called AtoMx, which is cloud-based only, limiting the accessibility, especially for users wanting to use their own high-processing-cluster. Concerning the visualizer of the PhenoCycler and MACSima, they are specific to multiplex imaging, i.e. no transcript can be shown. Contrary to the other visualizers, Xenium Explorer can be both (i) downloaded freely and (ii) supports open file formats. This makes it a reliable choice for conversion from SpatialData. Also, it supports missing data, i.e. it will not crash when reading multiplex imaging data (from which no transcripts are available).

### 9.4 Visualization with the Xenium Explorer

After using Sopa, the files required by the Xenium Explorer are created. In particular, a file called “experiment.xenium” can be opened in the Xenium Explorer. The later software is freely available for both Windows and MacOS. Sopa has been tested on versions 1.2 and 1.3 of the Xenium Explorer. We show two examples of visualization in Figure 1 (Xenium dataset, 10X) and Figure 2 (MERSCOPE dataset, Vizgen).

### 9.5 Image alignment with the Xenium Explorer

One challenge for spatial transcriptomics can be to align images from different technologies when they are run on the same sample. Most of the time, a simple affine transformation is enough to align them. Since Sopa create outputs in the Xenium Explorer, it is possible to use the alignment tool available on the software. It consists of applying some mirroring transformations, rotations, and alignment based on user-defined reference points. Then, the transformation matrix can be saved via the visualizer, which will create a “matrix.csv” transformation file. Afterwards, we can use this transformation matrix to align the new image on our Spatial-Data object and perform any operation available in Sopa. This can be done via the Sopa CLI, by specifying `sopa explorer add-aligned <sdata path> <image_path> <matrix_path>`. Typically, when adding an IF image, we can compute the mean channel intensity for all cells and for all channels.

**Supplementary Figure 1:**
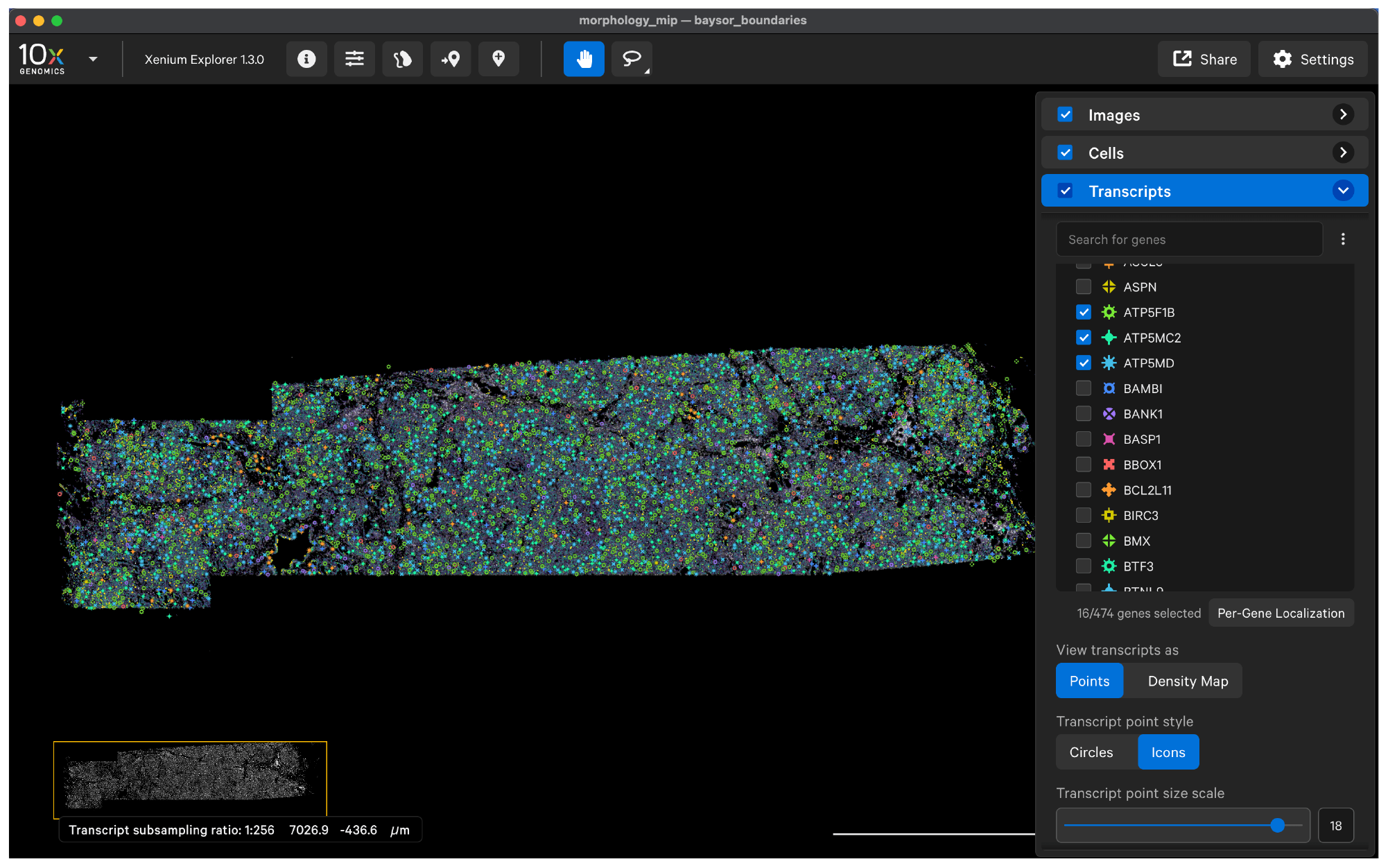
Xenium pancreas dataset (10X Genomics) open in the Xenium Explorer. The transcript panel is shown, with a few genes selected. Cells are coloured by a colour gradient representing transcript count.

### 9.6 Synthetic dataset generation

In order to demonstrate Sopa’s efficiency on multiple dataset sizes, we created synthetic datasets. Let *L* be the width of the image, and *d* be the cell density in the image. An evenly distributed grid of size 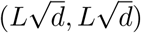 is generated, each vertex corresponding to a cell location. We apply a Gaussian noise of standard-deviation 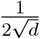 on these cell locations to have a more natural distribution of cells. Images are generated by applying a Gaussian blur of standard deviation 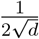 on the pixels at the location of the cell vertices, and 100 transcripts per cell are generated via a 2D Gaussian distribution of the same standard deviation.

**Supplementary Figure 2:**
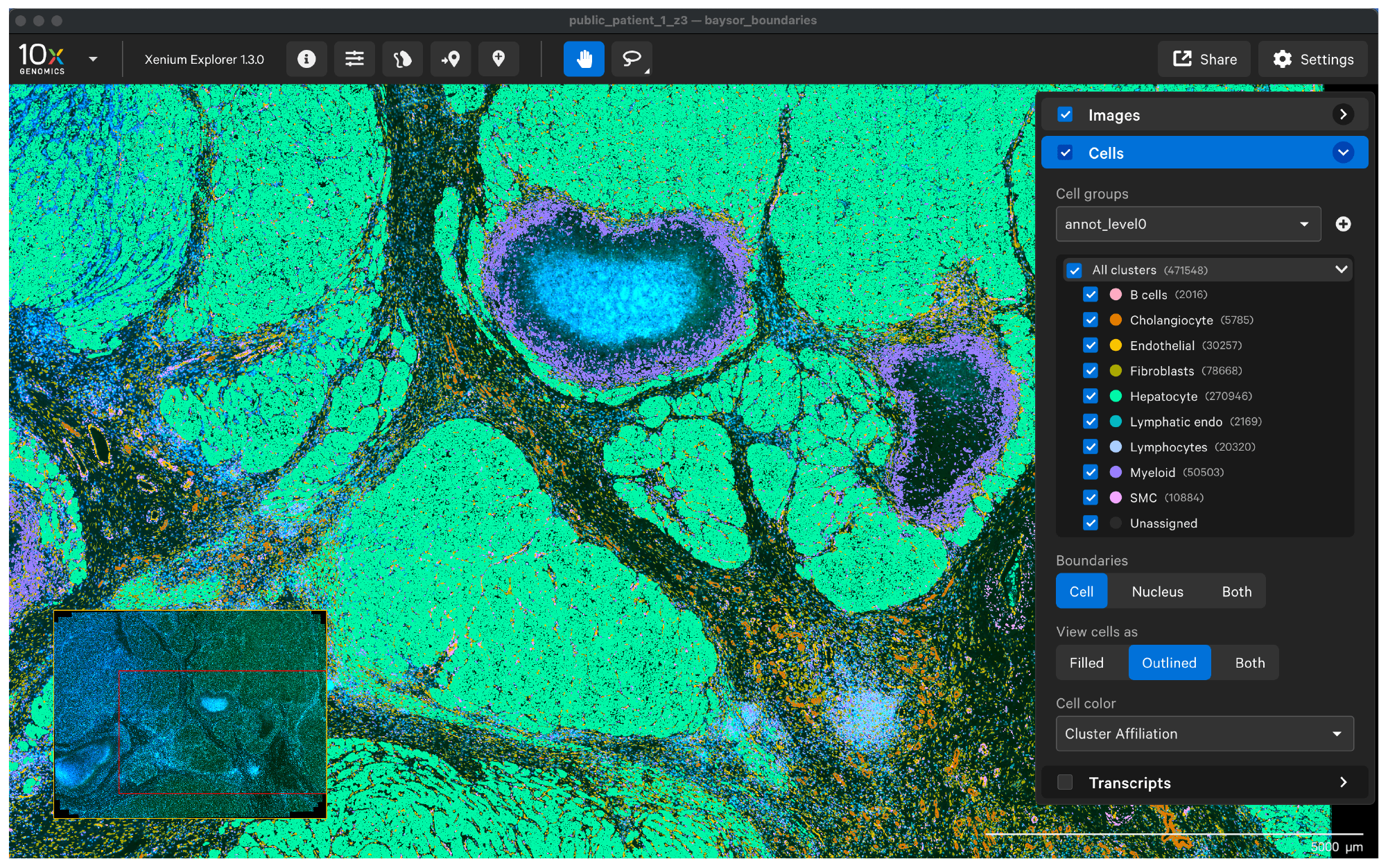
MERSCOPE liver dataset (Vizgen) open in the Xenium Explorer. The cell panel is shown, and the “*annot_level0*” category is displayed. Colors correspond to a cell-type.

### 9.7 Annotation of example datasets

Dataset annotation followed the procedure outlined in subsection 3.7. Automatic annotation utilized the following references: Liver dataset^6^ and Pancreas dataset^7^. Initial global annotation involved combining major cell populations, followed by refinement using Leiden clustering ^38^. Subsequent in-depth analysis employed manual annotation with Leiden clustering. For MACSima and PhenoCycler datasets, exclusion criteria involved DAPI, boundary staining, and low-quality proteins to enhance resolution. Manual clustering with Leiden was then applied for population annotation. Niche calculations were performed using STAGATE^27^. Niches were annotated based on cell type abundance and tissue structure, validated by a pathologist.

**Supplementary Figure 3:**
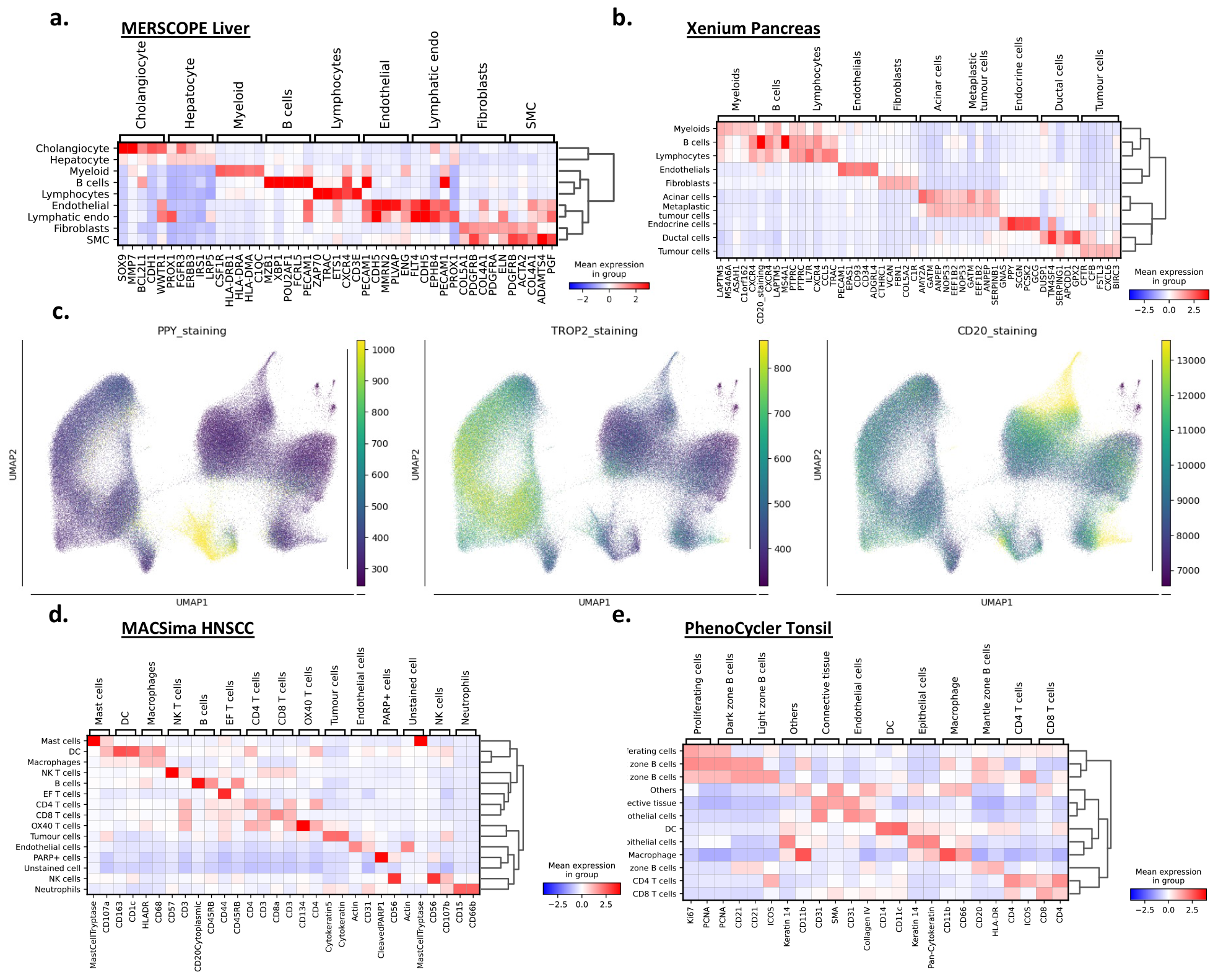
Visual validation of the annotations. **a**. Heatmap of genes expression per population on the MERSCOPE liver dataset. **b**. Heatmap of genes expression per population on the Xenium pancreas dataset. **c**. Protein staining per cell on the Xenium pancreas dataset after aligning the staining image to the original Xenium image. **d**. Heatmap of genes expression per population on the MACSima HNSCC dataset. **e**. Heatmap of genes expression per population on the Phenocycler tonsil dataset.

**Supplementary Figure 4:**
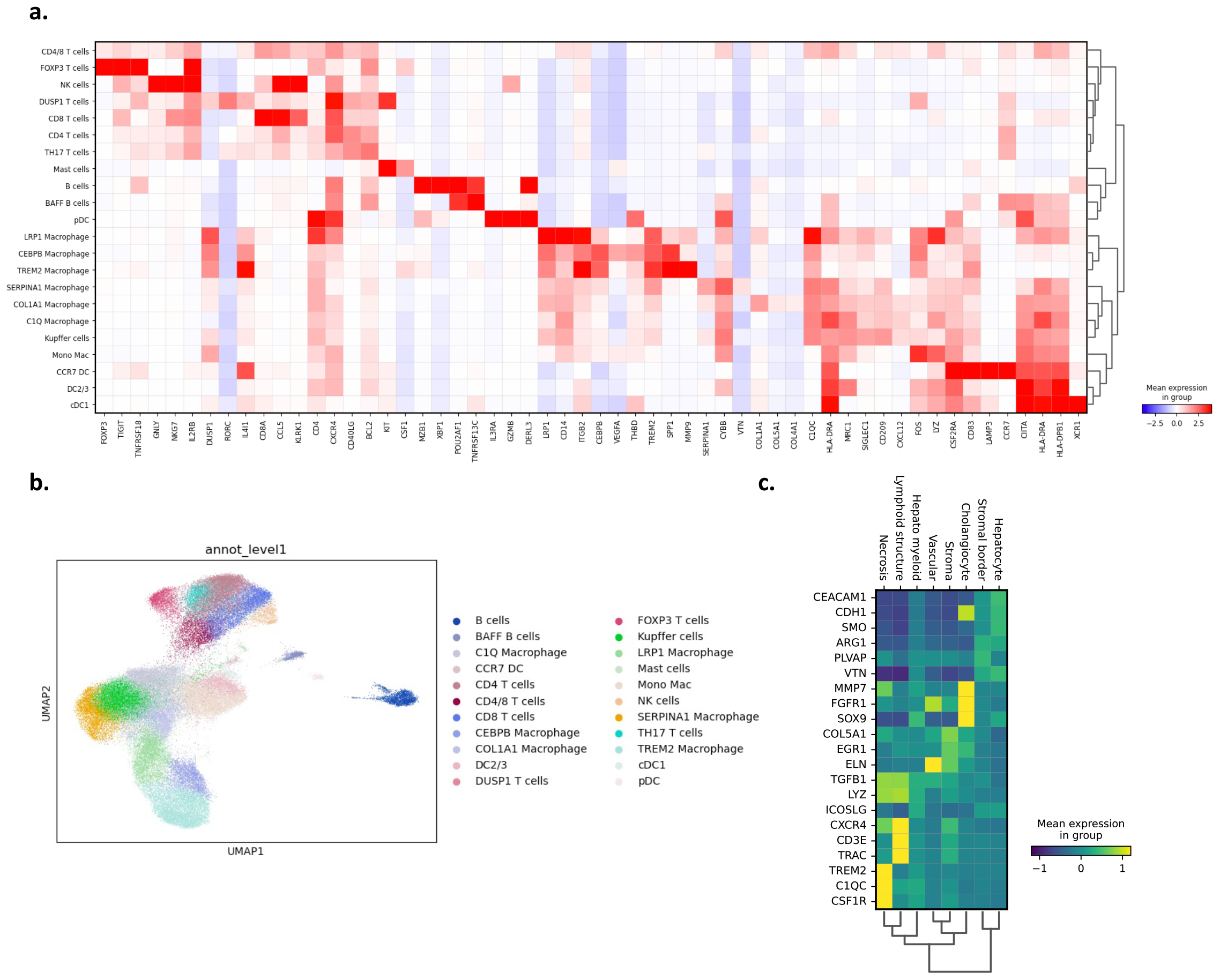
Annotation of MERSCOPE liver immune cells and niche differential gene expressions (DEGs). **a**. Heatmap of DEGs per immune population on the MERSCOPE liver dataset. **b**. UMAP of immune cells of the MERSCOPE liver dataset **c**. Heatmap of DEGs per niche of the MERSCOPE liver dataset.

## Footnotes

1 https://github.com/gustaveroussy/sopa

2 See subsection 3.1 and supplementary subsection 9.7 for more details

3 https://vizgen.github.io/vizgen-postprocessing/

4 https://www.10xgenomics.com/support/software/xenium-explorer/tutorials/xe-image-file-conversion

5 https://zarr.readthedocs.io/en/stable/index.html

6 https://www.immunesinglecell.org/atlas/liver

7 https://www.immunesinglecell.org/atlas/pancreas

## References

1. Bressan, D., Battistoni, G. & Hannon, G. J. The dawn of spatial omics. Science 381, eabq4964 (2023).

2. Rao, A., Barkley, D., Franca, G. S. & Yanai, I. Exploring tissue architecture using spatial transcriptomics. Nature 596, 211–220 (2021).

3. Moses, L. & Pachter, L. Museum of spatial transcriptomics. Nat Methods 19, 534–546 (2022).

4. Lewis, S. M. et al. Spatial omics and multiplexed imaging to explore cancer biology. Nat Methods 18, 997–1012 (2021).

5. Chen, K. H., Boettiger, A. N., Moffitt, J. R., Wang, S. & Zhuang, X. Spatially resolved, highly multiplexed RNA profiling in single cells. Science 348, aaa6090 (2015).

6. He, S. et al. High-plex imaging of RNA and proteins at subcellular resolution in fixed tissue by spatial molecular imaging. Nat Biotechnol 40, 1794–1806 (2022).

7. Kinkhabwala, A. et al. MACSima imaging cyclic staining (MICS) technology reveals combinatorial target pairs for CAR T cell treatment of solid tumors. Sci Rep 12, 1911 (2022).

8. Jhaveri, N. et al. Single-cell Spatial Metabolic and Immune Phenotyping of Head and Neck Cancer Tissues Identifies Tissue Signatures of Response and Resistance to Immunotherapy. 2023.05.30.540859 Preprint at 10.1101/2023.05.30.540859 (2023).

9. Chang, Q. et al. Imaging Mass Cytometry. Cytometry Part A 91, 160–169 (2017).

10. Spatial Gene and Protein Expression. 10x Genomics https://www.10xgenomics.com/products/spatialgene-and-protein-expression.

11. Jin, L. & Lloyd, R. V. In situ hybridization: Methods and applications. Journal of Clinical Laboratory Analysis 11, 2–9 (1997).

12. Merritt, C. R. et al. Multiplex digital spatial profiling of proteins and RNA in fixed tissue. Nat Biotechnol 38, 586–599 (2020).

13. Kumar, T. et al. A spatially resolved single-cell genomic atlas of the adult human breast. Nature 620, 181–191 (2023).

14. Chu, Y. et al. Pan-cancer T cell atlas links a cellular stress response state to immunotherapy resistance. Nat Med 29, 1550–1562 (2023).

15. Atta, L. & Fan, J. Computational challenges and opportunities in spatially resolved transcriptomic data analysis. Nat Commun 12, 5283 (2021).

16. Zeng, Z., Li, Y., Li, Y. & Luo, Y. Statistical and machine learning methods for spatially resolved transcriptomics data analysis. Genome Biology 23, 83 (2022).

17. Vandereyken, K., Sifrim, A., Thienpont, B. & Voet, T. Methods and applications for single-cell and spatial multi-omics. Nat Rev Genet 24, 494–515 (2023).

18. Dries, R. et al. Advances in spatial transcriptomic data analysis. Genome Res 31, 1706–1718 (2021).

19. Stringer, C., Wang, T., Michaelos, M. & Pachitariu, M. Cellpose: a generalist algorithm for cellular segmentation. Nat Methods 18, 100–106 (2021).

20. Petukhov, V. et al. Cell segmentation in imaging-based spatial transcriptomics. Nat Biotechnol 40, 345–354 (2022).

21. Biancalani, T. et al. Deep learning and alignment of spatially resolved single-cell transcriptomes with Tangram. Nat Methods 18, 1352–1362 (2021).

22. Palla, G. et al. Squidpy: a scalable framework for spatial omics analysis. Nat Methods 19, 171–178 (2022).

23. Hao, Y. et al. Dictionary learning for integrative, multimodal and scalable single-cell analysis. Nat Biotechnol 1–12 (2023).

24. Axelrod, S. et al. starfish: scalable pipelines for image-based transcriptomics. Journal of Open Source Software 6, 2440 (2021).

25. Cisar, C., Keener, N., Ruffalo, M. & Paten, B. A unified pipeline for FISH spatial transcriptomics. Cell Genomics 3, (2023).

26. Koster et al. Snakemake—a scalable bioinformatics workflow engine. Bioinformatics (2012).

27. Dong, K. & Zhang, S. Deciphering spatial domains from spatially resolved transcriptomics with an adaptive graph attention auto-encoder. Nat Commun 13, 1739 (2022).

28. McInnes, L., Healy, J. & Melville, J. UMAP: Uniform Manifold Approximation and Projection for Dimension Reduction. 1802.03426 [cs, stat] (2020).

29. Marconato, L. et al. SpatialData: an open and universal data framework for spatial omics. Preprint (2023).

30. Gillies, Sean et al., Manipulation and analysis of geometric objects in the Cartesian plane. Dask Development Team (2016).

31. Dask: Library for dynamic task scheduling.

32. geopandas/geopandas: v0.8.1. doi:10.5281/zenodo.3946761.

33. Wolf, F. A., Angerer, P. & Theis, F. J. SCANPY: large-scale single-cell gene expression data analysis. Genome Biology 19, 15 (2018).

34. Hoyer, S. & Hamman, J. xarray: N-D labeled Arrays and Datasets in Python. 5, 10 (2017).

35. McKinney, W. Data Structures for Statistical Computing in Python. in 56–61 (2010).

36. Barrett, P., Hunter, J., Miller, J. T., Hsu, J.-C. & Greenfield, P. matplotlib – A Portable Python Plotting Package. in (2005).

37. napari contributors (2019). napari: a multi-dimensional image viewer for python

38. Traag, V. A., Waltman, L. & van Eck, N. J. From Louvain to Leiden: guaranteeing well-connected communities. Sci Rep 9, 5233 (2019).

39. Virtanen, P. et al. SciPy 1.0: fundamental algorithms for scientific computing in Python. Nat Methods 17, 261–272 (2020).

40. Pedregosa, F. et al. Scikit-learn: Machine Learning in Python. J. Mach. Learn. Res. 12, 2825–2830 (2011).

41. Brodersen, P. Netgraph: Publication-quality Network Visualisations in Python. Journal of Open Source Software 8, 5372 (2023).

42. Wu, Z. et al. Graph deep learning for the characterization of tumour microenvironments from spatial protein profiles in tissue specimens. Nat. Biomed. Eng 6, 1435–1448 (2022).

43. Moore, J. et al. OME-NGFF: a next-generation file format for expanding bioimaging data-access strategies. Nat Methods 18, 1496–1498 (2021).

44. Virshup, I. et al. The scverse project provides a computational ecosystem for single-cell omics data analysis. Nat Biotechnol 41, 604–606 (2023).

45. Mulder, K. et al. Cross-tissue single-cell landscape of human monocytes and macrophages in health and disease. Immunity 54, 1883-1900.e5 (2021).

46. Khantakova, D., Brioschi, S. & Molgora, M. Exploring the Impact of TREM2 in Tumor-Associated Macrophages. Vaccines (Basel) 10, 943 (2022).

47. Sautes-Fridman, C., Petitprez, F., Calderaro, J. & Fridman, W. H. Tertiary lymphoid structures in the era of cancer immunotherapy. Nat Rev Cancer 19, 307–325 (2019).

48. Sharma, A. et al. Onco-fetal Reprogramming of Endothelial Cells Drives Immunosuppressive Macrophages in Hepatocellular Carcinoma. Cell 183, 377-394.e21 (2020).

49. Zhou, L. et al. Integrated Analysis Highlights the Immunosuppressive Role of TREM2+ Macrophages in Hepatocellular Carcinoma. Frontiers in Immunology 13, (2022).

50. Binnewies, M. et al. Targeting TREM2 on tumor-associated macrophages enhances immunotherapy. Cell Reports 37, 109844 (2021).

51. Molgora, M. et al. TREM2 Modulation Remodels the Tumor Myeloid Landscape, Enhancing Anti-PD-1 Immunotherapy. Cell 182, 886-900.e17 (2020).

52. Bijelic, L. & Rubio, E. R. Tumor Necrosis in Hepatocellular Carcinoma—Unfairly Overlooked? Ann Surg Oncol 28, 600–601 (2021).

53. Wei, T. et al. Tumor Necrosis Impacts Prognosis of Patients Undergoing Curative-Intent Hepatocellular Carcinoma. Ann Surg Oncol 28, 797–805 (2021).

54. Jass, J. R. Classification of colorectal cancer based on correlation of clinical, morphological and molecular features. Histopathology 50, 113–130 (2007).

55. Sharma, S., Sharma, M. C. & Sarkar, C. Morphology of angiogenesis in human cancer: a conceptual overview, histoprognostic perspective and significance of neoangiogenesis. Histopathology 46, 481–489 (2005).

